# Enhanced TRPV1 activation through TLR-4 and PKA signaling in Dorsal Root Ganglia Neurons

**DOI:** 10.64898/2026.06.24.734307

**Authors:** Julia Borges Paes Lemes, Kauê Franco Malange, Alisa Panichkina, Juliana Navia-Pelaez, Soo-Ho Choi, Maksim Dolmat, Gilson Gonçalves dos Santos, Sara Dochnal, Maripat Corr, Yury I. Miller, Tony Yaksh

**Affiliations:** Department of Anesthesiology, University of California, San Diego, CA, USA; Department of Medicine, University of California, San Diego, CA, USA; Department of Chemical and Nano Engineering, University of California, San Diego, CA, USA

**Keywords:** DRG, nociceptors, capsaicin, LPS, TLR-4, PKA, TRPV1

## Abstract

The excitability of afferents involved in nociceptive signaling reflects the interaction of several co-expressed membrane receptors. Current studies have shown that Toll-like receptor-4 (TLR-4) signaling can exacerbate excitation evoked by transient receptor potential vanilloid type 1 (TRPV1) activity, and this interaction plays a key role in driving and sustaining facilitated pain states. The mechanism by which this potentiated TRPV1 activity secondary to TLR-4 agonism occurs in sensory neurons remains unknown, although intracellular kinase activity is a strong candidate. To address this hypothesized linkage, neuronal cell cultures prepared from dorsal root ganglia (DRG) of male wildtype (WT) and *Tlr4^-/-^* mice were used to evaluate calcium transients of neurons after capsaicin administration in culture, pre-treated for 30 minutes with the TLR-4 agonist, lipopolysaccharide (LPS). TRPV1 protein expression at the neuron surface in cultured DRG cells with or without LPS treatment was quantified by flow cytometry assay. The roles of protein kinase A (PKA) and C were assessed using selective inhibitors (KT5720 for PKA and Chelerythrine chloride for PKC) applied to WT-DRG neurons or administered in vivo by intraplantar or intrathecal injection, prior to LPS and capsaicin administration. Behavioral effects of in vivo TRPV1 activation were assessed through paw flinch responses evoked by intraplantar capsaicin injection and by hind paw tactile thresholds measured by von Frey filaments. LPS incubation in cultured DRG neurons enhances the intensity of calcium influx following TRPV1 activation in WT but not *Tlr4^-/^* cells. The augmented calcium influx evoked by capsaicin was prevented by the inhibition of PKA but not PKC. Similarly, mice treated with LPS in the hind paw displayed greater nociceptive responding after capsaicin and increased tactile allodynia. The facilitated component was prevented by the local pre-treatment with the PKA inhibitor. Correspondingly, lumbar spinal blockade of PKA resulted in temporary reversal of hyperalgesia induced by intrathecal LPS injection in mice. Together, these results demonstrate the relevance of TLR-4 in modulating the excitability of nociceptor signaling by regulating TRPV1, thereby influencing pain transmission through PKA signaling.

## 1. INTRODUCTION

The transient receptor potential vanilloid type 1 (TRPV1) is a pain signal integrator that acts in both physiological and pathological conditions. TRPV1 activation leads to a conformational change in the channel, allowing the passage of cations, especially calcium (Ca^2+^), which induces membrane depolarization and action potential discharges, resulting in neuronal afferent activation [1]. Studies reported that TRPV1 initiated signaling can be substantially augmented by mediators released within the injury and inflammation site, affecting the electrical membrane thresholds of nociceptor soma and terminals. Such enhanced signaling contributes to the facilitated pain behavior phenotype, such as observed in arthritis [2] and fibromyalgia pathologic conditions [3].

A crucial principle is that neuronal channels and receptors coexist within an organizational matrix, the lipid raft, which permits interactions between these signaling components, leading to functional changes in indices of excitability [4]. One particularly important manifestation of this interaction occurs between TLR-4 and TRPV1 in the primary afferent neuron during neuropathic pain induced by a chemotherapy drug [4]. Activation of the TLR-4 expressed in lipid rafts results in increased proximity of TRPV1 receptors in neurons of murine DRG during peripheral neuropathic pain phenotype induced by paclitaxel [4]. In addition, TLR-4 activation by the agonist LPS, in HEK cells or trigeminal neurons, markedly potentiated Calcitonin Gene Related Peptide (CGRP) release upon TRPV1 activation by capsaicin [5–7]. Although the relationship between these two receptors has been studied, the intracellular molecular mechanism by which TRPV1 becomes sensitized via TLR-4 in nociceptors has not been characterized.

During tissue injury and inflammation, various inflammatory mediators activate multiple protein kinases that, in turn, phosphorylate TRPV1 [8,9], enhancing the channel’s responsiveness [10]. As LPS leads to the activation of Protein kinase (PK) A and C, we sought in the present work to determine if this linkage may mediate the augmentation by TLR-4 of activation of TRPV1 of the nociceptor soma terminal, and that inhibition of those kinases blocks TLR-4-initiated facilitation of TRPV1 activation. Here, we showed that PKA but not PKC participates in the augmented TRPV1 activity by increasing Ca^2+^ influx and protein phosphorylation after short-term treatment with LPS in DRG neurons. Comparable results were observed in vivo on capsaicin-evoked pain behavior.

## 2. METHODS

### 2.1. Animals

Adult male C57BL/6J mice or *Tlr4^-/-^* (8 weeks) mice were housed in isolated, ventilated racks, in a 12:12-hour light or dark cycle with a controlled temperature (22 °C). Food and water were provided ad libitum. The *Tlr4^-/-^* mice were a gift from Dr. S. Akira (Osaka University, Japan) and were backcrossed for ten generations onto the C57Bl/6 background [11]. Behavioral experiments followed the International Association for the Study of Pain (IASP) guidelines, and all animal procedures were approved by the UCSD IACUC (protocol number #S00137M).

### 2.2. Calcium imaging

Male mice were euthanized under 5% Isoflurane, followed by decapitation. Thoracic and lumbar DRGs were harvested and subjected to enzymatic dissociation, 1-hour incubation in 2% collagenase type IV (Gibco, # 17104019 Waltham, MA) solution, and 10 minutes in 0.25% trypsin (Gibco, 25200056) solution, at 36°C. Posterior, mechanical dissociation of the tissue was performed to obtain the isolated cells. Using the gradient method with 10% BSA (Miltenyi Biotec #130-091-376), neurons were separated from debris, kept in the BSA layer, while neuronal cells were deposited into the pellet. The cells were resuspended in DMEM supplemented with 0.5% FBS (media) and plated in a glass-bottom individual coverslip coated with Poly-DL-ornithine hydrobromide (Sigma P8638, St. Louis, MO). Cultures were maintained in the incubator at 37°C with a 5% CO_2_ atmosphere for 24 hours.

Cell cultures were loaded with the Ca^2+^ indicator Fluo-4 AM (5 mM; Invitrogen. #F23917, Waltham, MA) for 40 minutes at room temperature in HEPES-buffered saline (in mM): NaCl 122, KCl 3.3, CaCl_2_ 1.3, MgSO_4_ 0.4, KH_2_PO_4_ 1.2, HEPES 25, glucose 10, and adjusted with NaOH to pH 7.4. Coverslips were placed in a laminar flow perfusion chamber (Warner Instrument Corp, Holliston, MA) connected at one side with an input cannula that delivers the solutions and on the opposite side with a suction pump, to permit uninterrupted perfusion. The system contains three individual, separately valved reservoirs for different solutions.

Neuronal fluorescence intensity was recorded in an inverted Leica TCS SP5 confocal microscope. One field of view per coverslip was assessed, with images recorded at 63x/sec. In the standard study protocol, neurons were exposed to a buffered vehicle for 1 minute, then stimulated for 15 seconds with capsaicin (0.5 mM), buffered for 4 minutes, and finally stimulated with KCl (50 mM) for 5 seconds to confirm cellular viability. Neuronal Ca^2+^ responses were analyzed in 3 to 5 dishes from each group by selecting individual cells as ROIs (regions of interest) and calculating mean gray value variations on each cell using LAS AF version 2.7.3.9723 software. Data are presented as ΔF/F0, where F0 is the baseline, and the effect is quantified as maximal ΔF/F0.

### 2.3. Flow cytometry

DRGs were harvested and subjected to enzymatic dissociation using Neural Tissue Dissociation Kit (Miltenyi Biotec, San Diego, CA). After mechanical dissociation, neurons were resuspended in media and were equally separated into two groups, treated with either media or LPS (tlrl-3pelps, Envivogen®; 200 ng/ml; for 30 min;). After treatment, cells were fixed with PFA 4%, washed with PBS, and incubated for 45 min on ice with the following anti-bodies: APC-Cy7 live/dead Ghost dye (#18452, Cell Signaling, Danvers, MA), Anti-TRPV1 antibody (Novus biologics, #NBP199.427), and anti-Rabbit Alexa 488. Cells were analyzed using a CytoFLEX flow cytometer (Beckman Coulter, Brea, CA).

### 2.4. Drug injections

#### 2.4.1. Intrathecal

To perform intrathecal (i.t.) injections, mice were anesthetized using 3-5% isoflurane in oxygen and had the lower back shaved and disinfected. The L5 and L6 vertebrae were identified by palpation, and a 30G needle was inserted percutaneously into the midline between the L5 and L6 vertebrae [12]. A 5 μL injection of LPS (100 ng; tlrl-3pelps, Envivogen®), saline, or PKA inhibitor (PKAi; 10 µM; KT5720; Alomone Labs®) was administered following confirmation of successful entry in the intrathecal space, as indicated by a tail flick response [12].

#### 2.4.2. Subcutaneous

For subcutaneous (s.c.) injections, mice were gently restrained and drugs were administered into the right metatarsal footpad using an insulin syringe (30 G needle, BD Ultra-Fine®). Mice received a 20 μL injection of saline or PKA inhibitor (PKAi; 10 µM; KT5720; Alomone Labs®) followed by a 20 µL LPS injection (100 ng; tlrl-3pelps, Envivogen®). The mice were further assessed for pain behavior.

### 2.5. Pain behavior

#### 2.5.1. Assessment of Capsaicin-induced Nociceptive Response

Mice were inoculated with Capsaicin (5 µg/paw; M2028, Sigma Aldrich®) injected into the hindpaw as previously described on 2.4.1. The number of flinches were recorded immediately after capsaicin injection for up to 5 minutes. The nociceptive response was quantified as the total number of flinches and licks, with 3 seconds of licking considered equivalent to one flinch, as previously described [13].

#### 2.5.2. Assessment of paw mechanical thresholds

The tactile mechanical thresholds were assessed in the mice hind paw using the up-down method as previously described [14]. To measure the paw mechanical threshold, mice were placed individually in clear, plastic, bottomless cages over a wire mesh surface. Von Frey filaments (range from 2.44 to 4.31; 0.02-2.00 g) were applied to the plantar surface of both hind paws. Mechanical values for the left and right hind paws were measured and averaged to produce a single data point per assessment.

### 2.6. In vivo mechanistic studies

To assess the dependence of TLR-4-induced sensitization on PKA, mice received the PKA inhibitor 30 minutes before LPS administration via either the intrathecal or subcutaneous route. Tactile allodynia was evaluated at 3 and 24 hours following LPS injection. At the 24-hour time point, animals underwent the capsaicin test, as described in section 2.5.1, to determine the effects on TRPV1-mediated nociceptive responses.

### 2.7. Statistical Analysis

Data analysis was performed using GraphPad Prism v.10 software (GraphPad®, San Diego, USA). Normality of the values was tested using the D’Agostino-Pearson, Shapiro-Wilk, or Kolmogorov-Smirnov test, depending on the sample size. The T-Student test was used to compare two means. For comparisons involving more than two means, One-way or Two-way analysis of variance (ANOVA) was performed according to the experimental design. When significance indicated a statistical difference between means, the Tukey test was used to compare groups. The significance level adopted was p < 0.05.

## 3. RESULTS

### 3.1. TLR-4 agonist increases capsaicin-induced Ca^2+^ influx into sensory neurons

To evaluate the effect of LPS treatment on the intracellular Ca^2+^ levels of DRG neurons in response to capsaicin, cells were previously treated with media, LPS 100 ng/ml, or LPS 200 ng/ml for 30 minutes at 36 °C. Posteriorly, coverslips were washed out and incubated with the calcium indicator Fluo-4 AM (5 mM) for 40 minutes, followed by live confocal image recording (Figure 1A). As observed in Figure 1B, capsaicin administration evoked an immediate increase in intracellular Ca^2^ in the neurons as detected by increased fluorescence intensity. This change was detected in the media, LPS 100 ng/ml, and LPS 200 ng/ml treated groups (Figure 1B). However, pre-incubation with LPS at 200 ng/ml significantly augmented the neuronal response to capsaicin compared to the media or LPS at 100 ng/ml groups, as indicated by the maximum response analysis (Figure 1C). Suspended DRG neurons under LPS incubation (200 ng/ml for 30 minutes) resulted in significantly greater total TRPV1 protein surface expression when compared to cells in media, as detected by fluorescence analysis on flow cytometry (Figure 1D).

**Figure 1.**
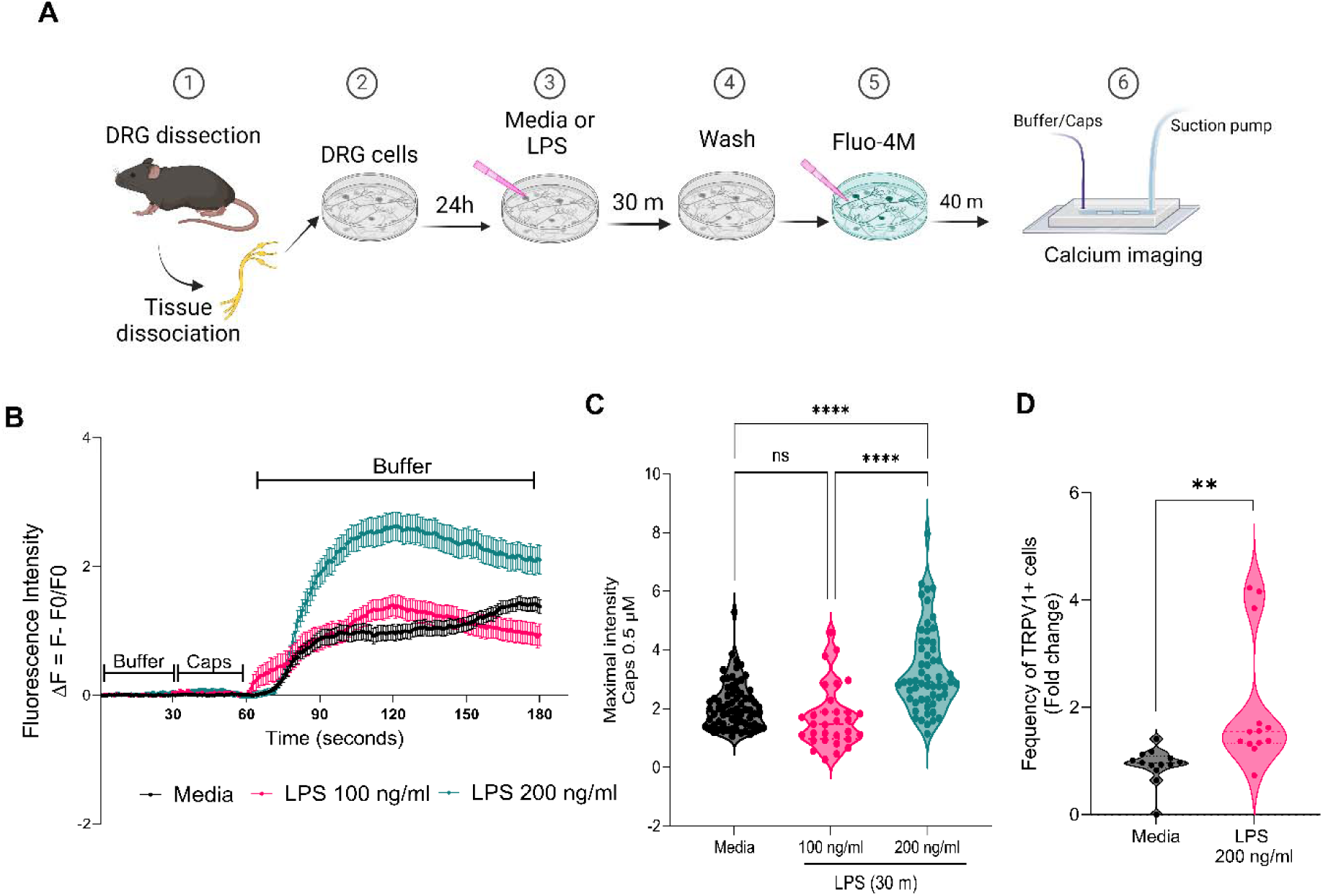
LPS increases capsaicin-induced Ca^2+^ influx in DRG neurons. (A) Experimental timeline: 1. Lumbar and thoracic DRG from naïve mice collected and dissociated; 2. Neurons were plated on individual glass coverslips for 24 hours; 3. Incubation of media or LPS 100-200 ng/ml for 30 minutes; 4. Single wash with buffer. 5. Calcium indicator incubation (Fluo-4 AM, 5 mM, 40 minutes); 6. Calcium imaging recording with capsaicin 0.5 µM stimulation for 30 seconds. (B) Fluorescence intensity (ΔF= F-F0/F0) of neurons after buffer and capsaicin, pretreated with media, LPS 100 or 200ng/ml. Data expressed as mean ± SEM. (C) The maximal intensity of response to the capsaicin stimuli. Five to nine neuronal culture dishes from 2 different wild-type mice were stimulated in vitro. Total number of cells recorded: media (n= 37), LPS 100 ng/ml (n= 56), LPS 200 ng/ml (n=31). (D) Flow cytometry: fold change of the frequency of TRPV1+ cells in media and LPS-treated group. Mean ± SEM; (C) One-way ANOVA, Tukey post-hoc test; (D) T-test **p< 0.001; ****p< 0.0001.

To confirm whether TLR-4 mediated the increased Ca^2^ influx in DRG neurons after LPS pre-treatment and triggered by capsaicin administration, a separate set of coverslips was used for TLR-4 antagonist (LPS-RS, 1 µg/µL) or vehicle (media) incubation for 30 minutes before LPS administration. Calcium imaging demonstrated that the TLR-4 antagonism reversed the enhanced fluorescence resulting from preconditioning DRG neurons with LPS (200 ng/ml) (Figure 2A). Similarly, DRG neurons from *Tlr4^-/-^* mice subjected to in vitro incubation with LPS (200 ng/ml, 30 minutes) did not present enhanced capsaicin-induced Ca^2^ influx compared to the control group (Figure 2B).

**Figure 2.**
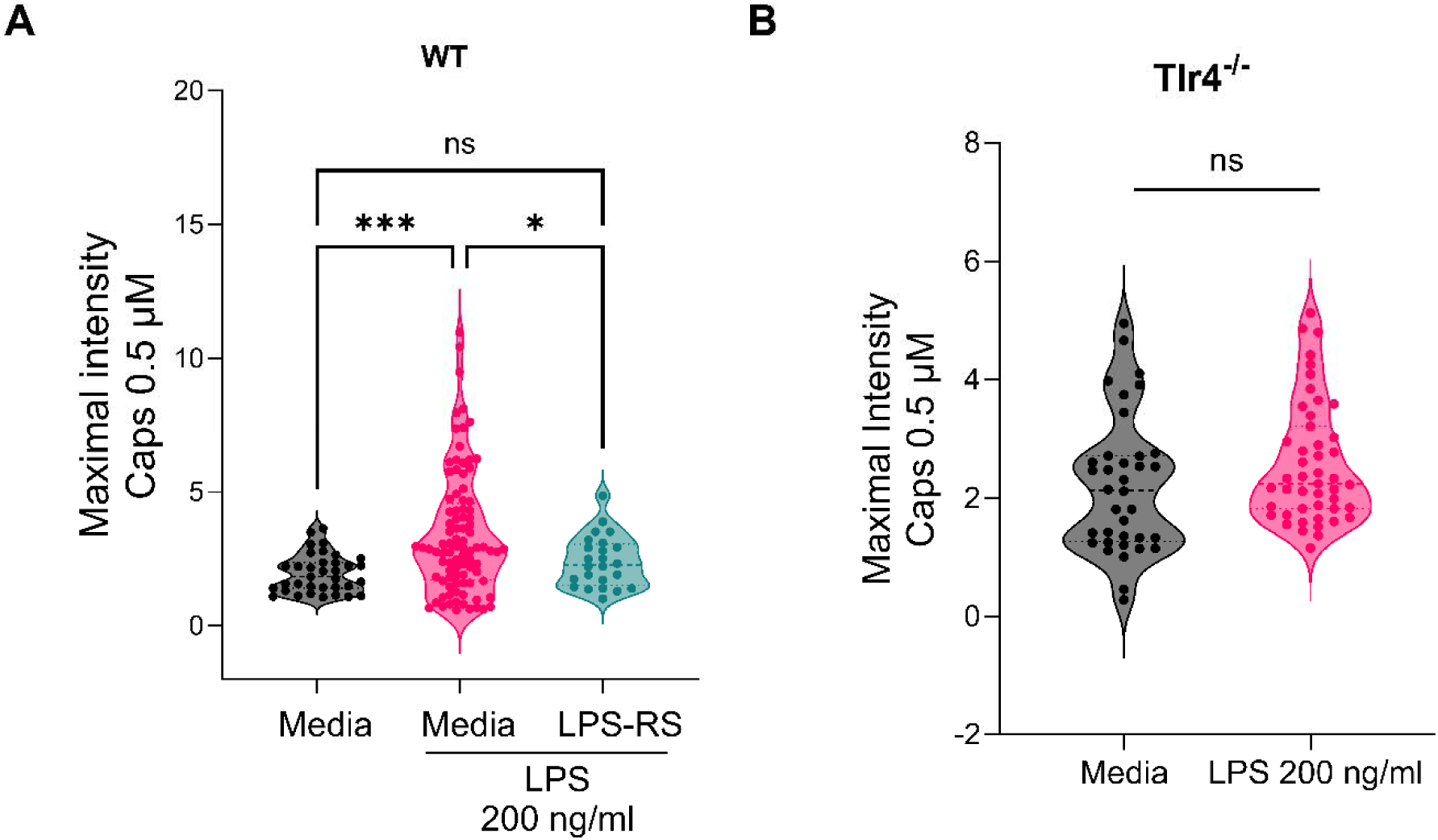
TLR-4 inhibition reversed neuronal TRPV1 response argumentation. (A) LPS-RS pre-treatment in DRG neurons reduced the maximal intensity of Ca^2+^ influx capsaicin-induced (0.5 µM) compared to the LPS group. Recorded cells: media n=35, LPS n=92, LPS-RS/LPS n=24. Mean ± SEM; One-way ANOVA, Tukey post-hoc test. (B) Maximal intensity of Tlr4-/- neurons treated with media or LPS (200 ng/ml) responded similarly to capsaicin stimuli (0.5 µM). Recorded cells: control n=36, LPS n=47. Mean ± SEM, T-test. *p < 0.05, ***p < 0.001.

### 3.2. Inhibition of PKA but not PKC reversed TRPV1-augmented activity induced by LPS

Next, selective inhibitors were applied to cultured DRG neurons to assess the role of PKC or PKA in modulating TRPV1 activity following LPS sensitization. Inhibitors PKCi (Chelerythrine chloride, 2 µM) or PKAi (KT5720, 0.3 µM) were diluted in media and applied to the neuronal cells for 30 minutes at 36 °C before LPS incubation (200 ng/ml, 30 minutes). All groups of cells, including control dishes, were subjected to a Ca^2^ influx assay induced by capsaicin (0.5 µM). The quantification of intracellular fluorescence revealed that neurons treated with LPS or PKCi + LPS exhibited a significantly higher intensity after capsaicin stimulation compared to the control group, whereas the cells pre-treated with PKAi showed a lower intensity (Figures 3A). The group treated with PKAi + LPS did not differ from the control, indicating that inhibition of PKA signaling reversed the augmented TRPV1 activity induced by LPS (Figures 3A).

**Figure 3.**
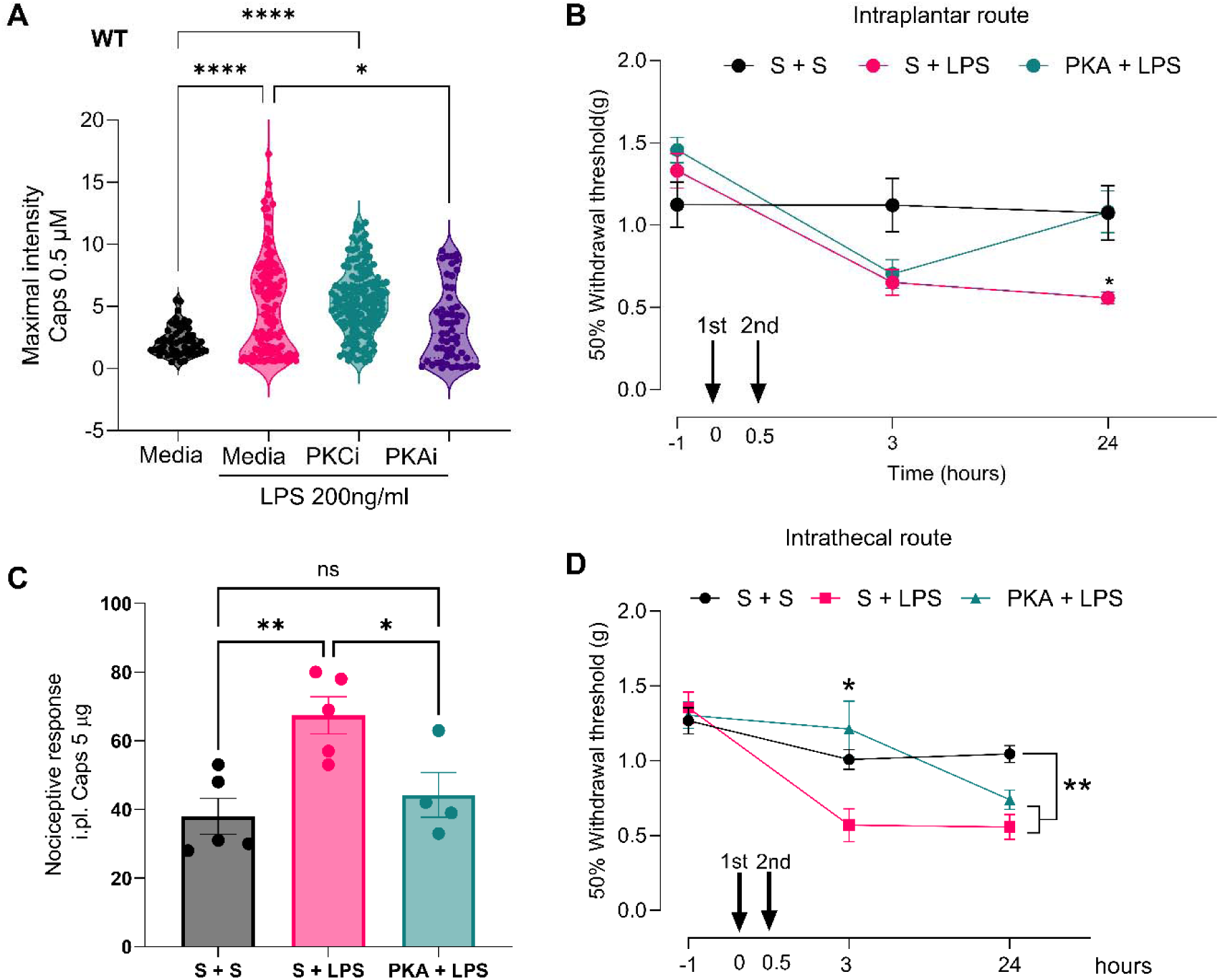
PKA inhibition diminishes the TRPV1-enhanced activity and peripheral pain induced by LPS. (A) Maximal intensity for DRG neurons, PKAi pre-treatment reduced the enhanced intensity of capsaicin-induced (0.5 µM) Ca^2+^ influx in neurons sensitized with LPS. Recorded cells: media n= 52, LPS n= 117, LPS-PKCi n= 146, LPA-PKAi n= 56. (B) Mechanical threshold in mice (n=7 mice per group) before and after intraplantar injection of Saline + Saline (S+S; 20 µL), Saline + LPS (S+LPS; LPS at 100 ng/20 µL), or PKAi +LPS (PKA + LPS; PKAi at 10 µM/20 µL; LPS at 100 ng/20 µL). (C) Nociceptive response after capsaicin (5 ug; 20 µL) injection to the hind paw. (D) Mechanical threshold in mice (n=7 mice per group) before and after intrathecal injection of S+S, S+LPS (LPS at 100 ng/5 µL), or PKAi +LPS (PKA + LPS; PKAi at 10 µM/5 µL; LPS at 100 ng/5 µL). Data represent the mean ± SEM; (A) and (C) One-way ANOVA. (B) and (D) Two-way ANOVA. Tukey post-hoc test. *p < 0.05, **p < 0.01, ****p < 0.0001.

Subsequently, to characterize the in vivo consequence of LPS at the peripheral nociceptor terminal, male mice received subcutaneous LPS injection (100 ng/paw) or saline (vehicle control) into the hind paw, and another group received PKAi (KT5720; 10 µM) 30 minutes before LPS injection. As observed in von Frey testing, groups treated with LPS displayed a significant decrease in mechanical threshold after 3 hours of injection, which persisted for 24h in the group saline + LPS (Figure 3B). In contrast, the group pre-treated locally with PKAi did not exhibit allodynia after 24h (Figure 2B). A similar reversal was observed in the capsaicin test performed 24h after intraplantar LPS injection; the saline + LPS group displayed higher number of paw flinches after capsaicin administration compared to the control group, and the PKAi + LPS group had a significant reduction (Figure 3C). Consistent with these peripheral observations, intrathecal administration of LPS also resulted in mechanical allodynia in the mice’s hind paw for 24h. This allodynia was also transiently reversed by pre-injection of PKAi intrathecally, at 3h, with recovery of allodynia at 24h, consistent with the transient effects of the inhibitor (Figure 3D).

## 4. DISCUSSION

The data presented in this work reveal that TRPV1 interacts functionally with TLR-4 in sensory neurons of the DRG, through PKA activation during sensitized states where TLR-4 is activated. This evidence was observed by in vitro Ca^2^ assay and behavioral hyperexcitability evaluation in mice.

Although no classic synapses occur in the sensory ganglia, a variety of neurotransmitters are locally released by non-neuronal cells (e.g., macrophages and glial cells) and by the afferent soma, making the DRG a site of effective integration of chemical signal transduction and augmenting nociceptive input [15]. Further, DRGs are located outside the blood-brain barrier, allowing circulating products such as large molecules and immune complexes to reach DRG cells, resulting in a local circuit driving nociceptor excitability and ectopic activity [15].

Thus, functional sensitization of afferent signaling is not limited to the peripheral terminal of the nociceptor, but such sensitization will also occur in the afferent soma. Nociceptive afferents are activated through specialized channels, such as TRPV1 [16], which undergo conformational changes in the presence of its agonist, leading to increased soma and terminal Ca^2+^ permeability [16]. The increase of Ca^2+^ in the cytosol is a result of extracellular Ca^2+^ or release from intracellular storage, such as mitochondria and endoplasmic reticulum (ER). For TRPV1, the majority Ca^2+^ increase is essentially from the extracellular medium and not from internal Ca^2+^ stores [16].

In nociceptors, the Ca^2+^ influx leads to i) acute depolarization resulting in activation of phosphorylating enzymes, local release of transmitters (substance P and CGRP), and ii) in the later periods, changes in protein expression [17]. Here, DRG-cultured neurons were loaded with a calcium indicator (Fluo-4 AM) and then subjected to a live calcium imaging assay using capsaicin as a stimulus. The control and treated dishes were first bathed in a continuous flow of buffer solution, which resulted in no changes in the intracellular fluorescence intensity, as expected. Brief exposure (30 seconds) to capsaicin (0.5 µM) resulted in an immediate Ca^2^ influx in the neurons, demonstrating soma activation. The group of cells treated with LPS at 200 ng/ml concentration presented higher intracellular Ca^2+^ influx after capsaicin stimuli than the control or LPS 100 ng/ml groups. These results are consistent with other reports, which show that acute treatment with LPS for 5 min led to an over two-fold increase in capsaicin-evoked currents in WT DRG neurons, which was absent in DRG neurons from the TLR-4-deficient mice [18]. Moreover, the co-localization of TLR-4 with TRPV1 receptors was demonstrated in a subpopulation of small-diameter neurons of DRG [19], and this interaction was suggested as a key factor in maintaining peripheral neuropathic pain induced by chemotherapy [4], arthritic pain [20], and hyperalgesia colitis-induced [18]. In our study, the use of the selective TLR-4 antagonist, LPS-RS, in WT-DRG neurons attenuated the facilitation of the capsaicin response induced by the TLR-4 agonist, LPS. Moreover, using DRG neurons from *Tlr4^-/-^*animals, we observed the absence of the augmented response to capsaicin after LPS incubation in WT DRGs. These findings confirmed that the TRPV1 sensitization induced by LPS occurs via TLR-4 receptors.

The acute augmented excitability observed may have arisen from increased cell surface expression of existing pools of TRPV1 protein and/or phosphorylation, enhancing channel functionality. The presence of increased cell surface TRPV1 protein was confirmed by flow cytometry analysis of cells treated with LPS at 200 ng/ml. Given the brief incubation time (30 min), the increase in TRPV1 protein detected is most likely related to protein trafficking of existing stores rather than de novo synthesis. For instance, in trigeminal cell culture, TNF-α incubation promoted an increase in surface expression of neuronal TRPV1 protein, leading to increased capsaicin-evoked Ca^2+^ influx. Interestingly, the total TRPV1 expression of the cells was not increased by western blot analysis after TNF-α stimulation; however, increased synaptic vesicle membrane protein (VAMP1) filled with TRPV1 was found close to the cell membrane [21].

TRPV1 receptors can be phosphorylated by several kinases, including PKA [22–24], PKC [25], calcium calmodulin–dependent kinase II (CaMKII) [26], or Src kinase [27]. The importance of residues Ser-116 and Thr-370 in the amino terminus is implicated in regulating TRPV1 excitability and desensitization [28]. Thus, seeking to understand the mechanism responsible for potentiated TRPV1 response following short-term TLR-4 agonism, protein kinase inhibitors (PKCi or PKAi) were applied to cultured DRG neurons before LPS incubation. As previously shown, the Ca^2^ influx induced by capsaicin in the presence of LPS was significantly higher than in the control group. However, PKA inhibition, but not PKC, reduced the sensitization of TRPV1 provoked by LPS. In contrast, in an osteoarthritic model induced by monoiodine acetate (MIA) in rats, pain-related behavior induced by intra-articular injection of capsaicin was inhibited by local pre-treatment with the PKC inhibitor [29]. In addition, phosphorylated-PKC (p-PKC ) increased and co-localized with TRPV1 in DRG neurons of MIA-OA rats [29]. Interestingly, protein levels of TRPV1 remained unchanged, but phosphorylated TRPV1 at Ser-800 increased in DRG neurons of MIA-OA rats [29].

Given the distinctness of issues related to neuronal-soma signaling and events occurring at the terminal of the axis, we sought to determine if the LPS-TLR-4 linkage evident in the cell body was recapitulated in the peripheral terminal. Mice treated with LPS (intraplantar route) displayed a greater number of paw flinches and mechanical allodynia after being injected with capsaicin, compared to the saline group. Local pre-treatment with PKAi into the paw reversed this effect, confirming the involvement of TRPV1 sensitization induced by LPS occurs via the TLR-4-PKA pathway, consistent with the data obtained in Ca^2^ assays. The behavioral response of flinching the paw after capsaicin administration is a motor reflex that occurs acutely (∼1-5 min) after activating TRPV1 receptors expressed on the nociceptive terminals (majority C-fiber type) [13,30]. The signal initiated at the peripheral terminal is transmitted by action potentials that reach the cell soma and then the central terminals at the dorsal root of the spinal cord. Interestingly, central administration of LPS to the lumbar spinal cord via intrathecal injection, a technique that also reaches the DRGs (L3-L5), resulted in a lower paw mechanical threshold, which was transiently attenuated by PKAi prior to LPS injection.

In summary, the present study has provided evidence that activation of TLR-4 in primary afferents at soma or terminals by LPS promotes increased TRPV1 function via PKA activation. These findings extend our understanding of pain mechanisms in the afferent signaling linkages and open doors for future studies to refine therapeutic approaches.

## 5. AUTHOR CONTRIBUTIONS

Study design, experiment execution, data analysis, and manuscript draft: J.B.P.L, Y.I.M., T.L.Y. Formal analysis, investigation, data curation: J.B.P.L., K.F.M., J.NP, G.G.S, A.P., S.D., M.D., MP.C, C. S-H. Writing—review and editing, visualization: J.B.P.L., K.F.M., A.P., S.D, MP.C, Y.I.M., T.L.Y. Supervising, conceptualizing, and project administration: Y.M., T.L.Y.

## 6. AKNOWLEDGMENTS

This was funded by NIH grant R01NS132483 given to TLY and Y.I.M.

## REFERENCES

1. Gao, N.; Li, M.; Wang, W.; Liu, Z.; Guo, Y. The dual role of TRPV1 in peripheral neuropathic pain: pain switches caused by its sensitization or desensitization. Front Mol Neurosci 2024, 17, 1400118, doi:10.3389/fnmol.2024.1400118.

2. Qu, Y.; Fu, Y.; Liu, Y.; Liu, C.; Xu, B.; Zhang, Q.; Jiang, P. The role of TRPV1 in RA pathogenesis: worthy of attention. Front Immunol 2023, 14, 1232013, doi:10.3389/fimmu.2023.1232013.

3. Yuksel, E.; Naziroglu, M.; Sahin, M.; Cig, B. Involvement of TRPM2 and TRPV1 channels on hyperalgesia, apoptosis and oxidative stress in rat fibromyalgia model: Protective role of selenium. Sci Rep 2017, 7, 17543, doi:10.1038/s41598-017-17715-1.

4. Navia-Pelaez, J.M.; Borges Paes Lemes, J.; Gonzalez, L.; Delay, L.; Dos Santos Aggum Capettini, L.; Lu, J.W.; Goncalves Dos Santos, G.; Gregus, A.M.; Dougherty, P.M.; Yaksh, T.L.; et al. AIBP regulates TRPV1 activation in chemotherapy-induced peripheral neuropathy by controlling lipid raft dynamics and proximity to TLR4 in dorsal root ganglion neurons. Pain 2023, 164, e274-e285, doi:10.1097/j.pain.0000000000002834.

5. Diogenes A, F.C., Akopian AN, Henry MA, Hargreaves KM. LPS Sensitizes TRPV1 via Activation of TLR4 in Trigeminal Sensory Neurons. Journal of Dental Research 2011, 759–764, doi:10.1177/0022034511400225.

6. Min, H.; Lee, H.; Lim, H.; Jang, Y.H.; Chung, S.J.; Lee, C.J.; Lee, S.J. TLR4 enhances histamine-mediated pruritus by potentiating TRPV1 activity. 2014, doi:10.1186/s13041-014-0059-9.

7. Min, H.; Cho, W.H.; Lee, H.; Choi, B.; Kim, Y.J.; Lee, H.K.; Joo, Y.; Jung, S.J.; Choi, S.Y.; Lee, S.;, et al. Association of TRPV1 and TLR4 through the TIR domain potentiates TRPV1 activity by blocking activation-induced desensitization. Mol Pain 2018, 14, 1744806918812636, doi:10.1177/1744806918812636.

8. Amadesi, S.; Cottrell, G.S.; Divino, L.; Chapman, K.; Grady, E.F.; Bautista, F.; Karanjia, R.; Barajas-Lopez, C.; Vanner, S.; Vergnolle, N.;, et al. Protease-activated receptor 2 sensitizes TRPV1 by protein kinase Cepsilon- and A-dependent mechanisms in rats and mice. J Physiol 2006, 575, 555–571, doi:10.1113/jphysiol.2006.111534.

9. Levine, J.D.; Alessandri-Haber, N. TRP channels: targets for the relief of pain. Biochim Biophys Acta 2007, 1772, 989–1003, doi:10.1016/j.bbadis.2007.01.008.

10. Joseph, J.; Qu, L.; Wang, S.; Kim, M.; Bennett, D.; Ro, J.; Caterina, M.J.; Chung, M.K. Phosphorylation of TRPV1 S801 Contributes to Modality-Specific Hyperalgesia in Mice. J Neurosci 2019, 39, 9954–9966, doi:10.1523/JNEUROSCI.1064-19.2019.

11. Hoshino, K.; Takeuchi, O.; Kawai, T.; Sanjo, H.; Ogawa, T.; Takeda, Y.; Takeda, K.; Akira, S. Cutting Edge: Toll-Like Receptor 4 (TLR4)-Deficient Mice Are Hyporesponsive to Lipopolysaccharide: Evidence for TLR4 as the Lps Gene Product. The Journal of Immunology 1999, 162, 3749–3752, doi:10.4049/jimmunol.162.7.3749.

12. Janice L.K. Hylden, G.L.W. Intrathecal morphine in mice: A new technique. European Journal of Pharmacology 1980, 67, 313–316, doi:10.1016/0014-2999(80)90515-4.

13. Lemes, J.B.P.; de Campos Lima, T.; Santos, D.O.; Neves, A.F.; de Oliveira, F.S.; Parada, C.A.; da Cruz Lotufo, C.M. Participation of satellite glial cells of the dorsal root ganglia in acute nociception. Neurosci Lett 2018, 676, 8–12, doi:10.1016/j.neulet.2018.04.003.

14. Chaplan, S.R.; Bach, F.W.; Pogrel, J.W.; Chung, J.M.; Yaksh, T.L. Quantitative assessment of tactile allodynia in the rat paw. Journal of Neuroscience Methods 1994, 53, 55–63, 10.1016/0165-0270(94)90144-9.

15. Berta, T.; Qadri, Y.; Tan, P.H.; Ji, R.R. Targeting dorsal root ganglia and primary sensory neurons for the treatment of chronic pain. Expert Opin Ther Targets 2017, 21, 695–703, doi:10.1080/14728222.2017.1328057.

16. Maximiano, T.K.E.; Carneiro, J.A.; Fattori, V.; Verri, W.A. TRPV1: Receptor structure, activation, modulation and role in neuro-immune interactions and pain. Cell Calcium 2024, 119, 102870, doi:10.1016/j.ceca.2024.102870.

17. Julius, D. TRP channels and pain. Annu Rev Cell Dev Biol 2013, 29, 355–384, doi:10.1146/annurev-cellbio-101011-155833.

18. Wu, Y.; Wang, Y.; Wang, J.; Fan, Q.; Zhu, J.; Yang, L.; Rong, W. TLR4 mediates upregulation and sensitization of TRPV1 in primary afferent neurons in 2,4,6-trinitrobenzene sulfate-induced colitis. Mol Pain 2019, 15, 1744806919830018, doi:10.1177/1744806919830018.

19. Li, Y.; Adamek, P.; Zhang, H.; Tatsui, C.E.; Rhines, L.D.; Mrozkova, P.; Li, Q.; Kosturakis, A.K.; Cassidy, R.M.; Harrison, D.S.;, et al. The Cancer Chemotherapeutic Paclitaxel Increases Human and Rodent Sensory Neuron Responses to TRPV1 by Activation of TLR4. J Neurosci 2015, 35, 13487–13500, doi:10.1523/JNEUROSCI.1956-15.2015.

20. Dos Santos, G.G.; Jimenez-Andrade, J.M.; Munoz-Islas, E.; Candanedo-Quiroz, M.E.; Cardenas, A.G.; Drummond, B.; Pham, P.; Stilson, G.; Hsu, C.C.; Delay, L.;, et al. Role of TLR4 activation and signaling in bone remodeling, and afferent sprouting in serum transfer arthritis. Arthritis Res Ther 2024, 26, 212, doi:10.1186/s13075-024-03424-4.

21. Meng, J.; Wang, J.; Steinhoff, M.; Dolly, J.O. TNFalpha induces co-trafficking of TRPV1/TRPA1 in VAMP1-containing vesicles to the plasmalemma via Munc18-1/syntaxin1/SNAP-25 mediated fusion. Sci Rep 2016, 6, 21226, doi:10.1038/srep21226.

22. Vlachová, V.; Teisinger, J.; Sušánková, K.; Lyfenko, A.; Ettrich, R.; Vyklický, L. Functional role of C-terminal cytoplasmic tail of rat vanilloid receptor 1. Journal of Neuroscience 2003, 23, 1340–1350.

23. De Petrocellis, L.; Harrison, S.; Bisogno, T.; Tognetto, M.; Brandi, I.; Smith, G.D.; Creminon, C.; Davis, J.B.; Geppetti, P.; Di Marzo, V. The vanilloid receptor (VR1)-mediated effects of anandamide are potently enhanced by the cAMP-dependent protein kinase. J Neurochem 2001, 77, 1660–1663, doi:10.1046/j.1471-4159.2001.00406.x.

24. Rathee, P.K.; Distler, C.; Obreja, O.; Neuhuber, W.; Wang, G.K.; Wang, S.-Y.; Nau, C.; Kress, M. PKA/AKAP/VR-1 module: A common link of Gs-mediated signaling to thermal hyperalgesia. Journal of Neuroscience 2002, 22, 4740–4745.

25. Bhave, G.; Hu, H.-J.; Glauner, K.S.; Zhu, W.; Wang, H.; Brasier, D.; Oxford, G.S.; Gereau IV, R.W. Protein kinase C phosphorylation sensitizes but does not activate the capsaicin receptor transient receptor potential vanilloid 1 (TRPV1). Proceedings of the national academy of sciences 2003, 100, 12480–12485.

26. Jung, J.; Shin, J.S.; Lee, S.Y.; Hwang, S.W.; Koo, J.; Cho, H.; Oh, U. Phosphorylation of vanilloid receptor 1 by Ca2+/calmodulin-dependent kinase II regulates its vanilloid binding. J Biol Chem 2004, 279, 7048-7054, doi:10.1074/jbc.M311448200.

27. Jin, X.; Morsy, N.; Winston, J.; Pasricha, P.J.; Garrett, K.; Akbarali, H.I. Modulation of TRPV1 by nonreceptor tyrosine kinase, c-Src kinase. American Journal of Physiology-Cell Physiology 2004, 287, C558–C563.

28. Mohr, D.C.; Hart, S.L.; Julian, L.; Cox, D.; Pelletier, D. Association between stressful life events and exacerbation in multiple sclerosis: a meta-analysis. Bmj 2004, 328, doi:10.1136/bmj.38041.724421.55.

29. Koda, K.; Hyakkoku, K.; Ogawa, K.; Takasu, K.; Imai, S.; Sakurai, Y.; Fujita, M.; Ono, H.; Yamamoto, M.; Fukuda, I.;, et al. Sensitization of TRPV1 by protein kinase C in rats with mono-iodoacetate-induced joint pain. Osteoarthritis Cartilage 2016, 24, 1254–1262, doi:10.1016/j.joca.2016.02.010.

30. Lemes, J.B.P.; Malange, K.F.; Carvalho, N.S.; Neves, A.F.; Urban-Maldonado, M.; Kempe, P.R.G.; Nishijima, C.M.; Fagundes, C.C.; Lotufo, C.; Suadicani, S.O.;, et al. Blocking Pannexin 1 Channels Alleviates Peripheral Inflammatory Pain but not Paclitaxel-Induced Neuropathy. J Integr Neurosci 2024, 23, 64, doi:10.31083/j.jin2303064.

